# Balancing Selection at *HLA-G* Modulates Fetal Survival, Preeclampsia and Human Birth Sex Ratio

**DOI:** 10.1101/851089

**Authors:** S. Wedenoja, M. Yoshihara, H. Teder, H. Sariola, M. Gissler, S. Katayama, J. Wedenoja, I.M. Häkkinen, S. Ezer, N. Linder, J. Lundin, T. Skoog, E. Sahlin, E. Iwarsson, K. Pettersson, E. Kajantie, M. Mokkonen, S. Heinonen, H. Laivuori, K. Krjutškov, J. Kere

**Author notes:** equal contributions (H. Laivuori, K. Krjutškov, J. Kere).

## Abstract

The population sex ratio is thought to be maintained through balancing selection on rare phenotypes. However, empirical evidence for genetic influence has thus far proven elusive. We combined 1000 Genomes data and large cohorts to study human sex ratios. We found underrepresentation of male offspring in preeclampsia, a serious pregnancy disorder with uncertain pathogenesis. Genetic variation of fetal *human leukocyte antigen G* (*HLA-G*), regulating maternal anti-fetal immune responses, was found to be under balancing selection. Sex-linked downregulation of *HLA-G* and upregulation of *interferon alpha-1* (*IFNA1*) expression contribute to loss of fetal immunotolerance in preeclampsia and suggest hydroxychloroquine as a treatment option. Our findings indicate that an evolutionary trade-off between fetal immunotolerance and protection against infections promotes genetic diversity in *HLA-G*, thereby maintaining human sex ratios.

**One Sentence Summary:** Fetal *HLA-G* modulates human sex ratio.

## Main Text

Whether human male-to-female sex ratios exhibit appreciable variation has been a subject of debate in the biological sciences. Recent large-scale data have confirmed an unbiased sex ratio at conception: equal numbers of X-bearing or Y-bearing sperm fertilize human eggs. However, the global birth sex ratio of 106 males for every 100 females, already seen in 20-week fetuses, indicates that more females are lost in early human pregnancies. In contrast, male fetuses might be at higher risk for late miscarriages and stillbirths *(1)*. Together, these findings suggest that undetermined fetal sex-specific mechanisms contribute to human pregnancy success and phenotypic variation in the sex ratio arises during pregnancy. Negative frequency-dependent selection has been proposed to keep sex ratios balanced *(2)*, however to our knowledge, there is a lack of empirical evidence across all species for a genetic link to sex ratio.

In humans, balancing selection at the human major histocompatibility (MHC) locus has been proposed to modulate a deficiency of human leukocyte antigen (HLA) homozygotes through maternal-fetal interaction *(3)*, without regard for sex-specificity. Increased HLA similarity in couples is associated with recurrent miscarriages, further supporting the need for the fetus to differ from maternal HLA *(4)*. Although the exact modifier locus remains unclear, only a limited pattern of HLA antigens (HLA-C, HLA-G, HLA-E, HLA-F) are expressed by fetal trophoblasts to prevent rejection by maternal immune cells *(5)*. Of these, HLA-G is the most studied due to its trophoblast-restricted expression and multiple isoforms inducing both local and systemic immunotolerance *(6)*.

Tens of studies, with still inconclusive results, have searched for the link between HLA-G, miscarriages, and the hypertensive pregnancy disorder preeclampsia *(7)*. While reduced HLA-G expression observed in the preeclamptic placenta is expected to facilitate maternal immune reactions to fetal alloantigens, its role in the pathogenesis is not yet clear *(8)*. Maternal immune maladaptation in preeclampsia, as one of the major theories, is supported by the shift from antibody-mediated T-helper 2 (Th2) and regulatory T cells (Treg) to cell-mediated T-helper 1 (Th1) responses, and by aberrant natural killer (NK) cell activity *(9, 10)*. This study was motivated by the proposed but still disputed sex bias in preeclamptic births *(11)*, and particularly, the low male/female ratio in children of both women and men descending from a proband with preeclampsia *(12)*. We hypothesized that male fetal loss reflects failure in fetal HLA-G mediated immunotolerance in preeclampsia, stronger maternal immune responses towards male fetuses, and demonstrates the particular role of HLA-G in maternal-fetal interaction that affects fetal survival in human pregnancy.

We examined birth sex ratios (male/female) in a Finnish population cohort of 1.79 million live and stillbirths. This cohort included 38,752 preeclamptic births (2.2%). The sex ratio was 1.02 in preeclamptic and 1.05 in non-preeclamptic births (P=0.006), and the earlier the gestational age at birth, the lower the number of male offspring in preeclampsia (Spearman correlation coefficient, 0.80; P=0.007) (Fig. 1A). The sex bias was in striking agreement with Norwegian data on 1.82 million births and 44,000 preeclamptic pregnancies *(13)*, and with others *(14)*. Together with the data on unbiased human sex ratio at conception *(1)* and familial association of preeclampsia and miscarriages *(15)*, our results suggested male fetuses being lost before the onset of early preeclampsia, or linked to a higher risk of term disease, or both.

**Fig. 1.**
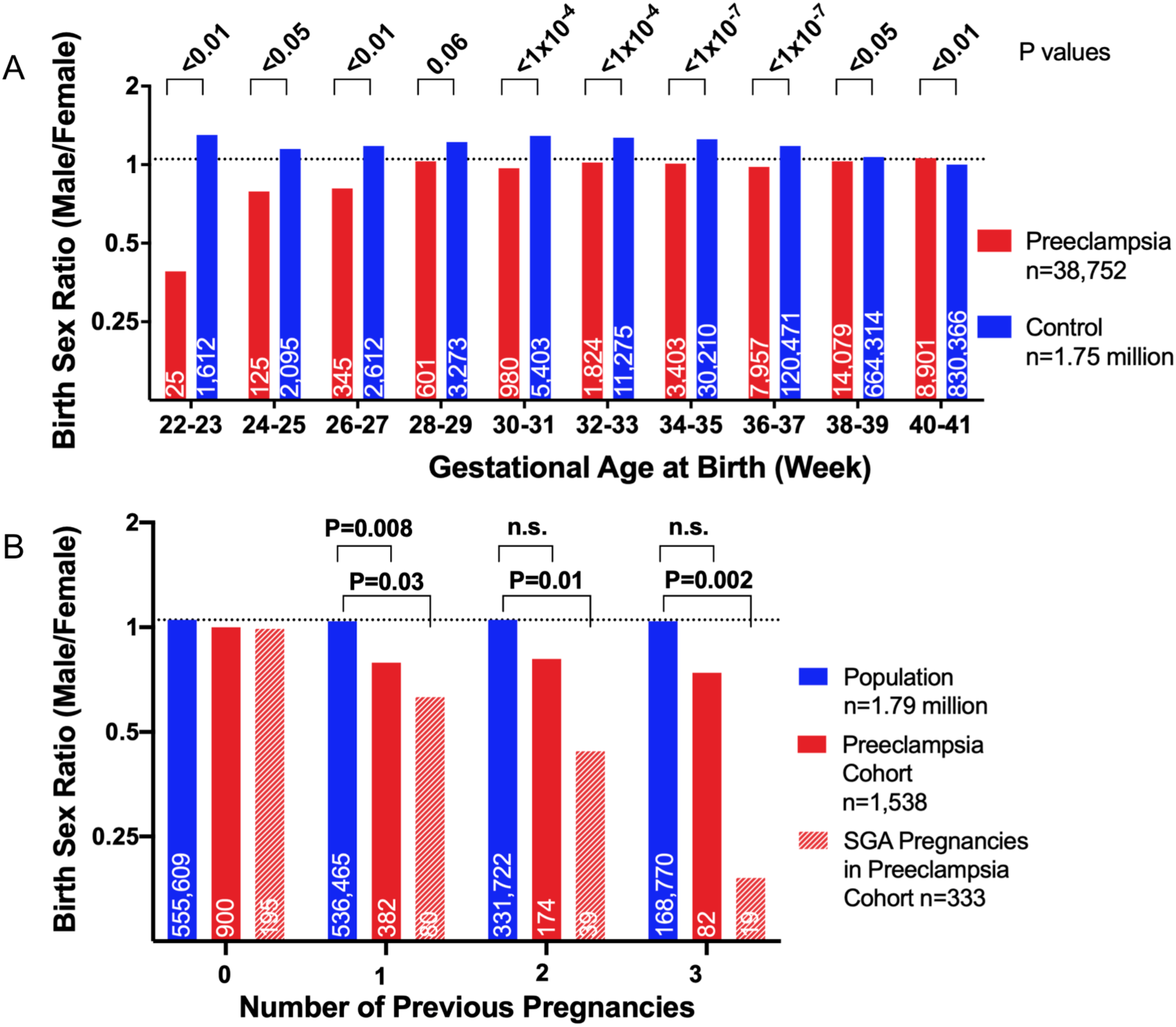
Biased Birth-Sex Ratio in Preeclampsia. (A) The change from the female biased birth sex ratio in early preeclampsia to male biased in term disease, and the opposite pattern from male biased in early to balanced sex ratio in term births without preeclampsia. Preeclamptic births after 41 weeks were rare (n=512) and the sex ratio of 0.99 was comparable with that of the controls (data not shown). (B) The relationship between the number of previous pregnancies and the birth sex ratios in the Finnish population, in preeclamptic pregnancies of the FINNPEC series, and in a subgroup of preeclamptic pregnancies with small for gestational age (SGA) offspring. Numbers of individuals in each group are represented in the bars and P values (Chi-square test) for birth sex differences above the graphs. Dashed lines indicate the population birth sex ratio of 1.05 and n.s. denotes non-significant.

We studied the possible association between male fetal loss and preeclampsia. Using population-level data on women with one or more miscarriages, we demonstrated the reduced sex ratio of 0.98 in consecutive preeclamptic (n=6,469) compared with 1.04 in non-preeclamptic births (n=355,683; P=0.03). In the FINNPEC preeclampsia cohort (n=1,538), the higher number of previous pregnancies was associated with reduced number of male offspring in preeclamptic women, especially in pregnancies with small for gestational age (SGA) offspring (Fig. 1B). We also studied a Swedish stillbirth cohort (n=277), which showed increased intrauterine loss of male fetuses with the sex ratio of 1.18. Furthermore, pregnancy complications were overrepresented in male offspring in the Finnish preeclampsia family series *(16)*. Altogether, our results suggest that maternal immune reactions to fetal alloantigens, and stronger responses to the male-specific Y chromosomal histocompatibility (H-Y) antigen *(17)*, contribute to fetal survival and preeclampsia pathogenesis.

*HLA-G* downregulation is linked to its 3’ untranslated regulatory region (3’UTR; Fig. S1) which is associated with distinct full-length haplotypes, and modulates mRNA stability and decay, microRNA targeting, and splicing *(6, 18)*. To uncover the link between *HLA-G* and the sex ratio, we studied *HLA-G* 3’UTR haplotype pairs, diplotypes, representing functional units. In the 1000 Genomes series (n=2504) *(19)*, sex ratios were modulated by the *HLA-G* 3’UTR diplotypes (Generalized linear model, GLM: F_21,24_=2.27, P=0.03; Fig. S2). Rare diplotypes showed opposite patterns of sex ratios in preeclampsia offspring (Fig. 2A) and stillborn fetuses (Fig. 2B). The distribution of *HLA-G* 3’UTR diplotypes and the sex ratio differed between offspring from preeclamptic (Fig. 2A) and control pregnancies (Fig. 2C) in the FINNPEC series (GLM: F_13,21_=9.15, P<0.001). We found a similar tendency in Africans versus Non-Africans in the 1000 Genomes data (GLM: F_1,24_=8.47, P=0.008; Fig. S2), the Africans having the highest global prevalence of preeclampsia *(20)*. Altogether, rare diplotypes showed more variation in the sex ratio than the common ones, which were close to the balance of 1.0. These results support a pattern of negative frequency-dependent selection, a form of balancing selection in which rare alleles are favored and the fitness of each allele decreases as its frequency increases *(21-23)*.

**Fig. 2.**
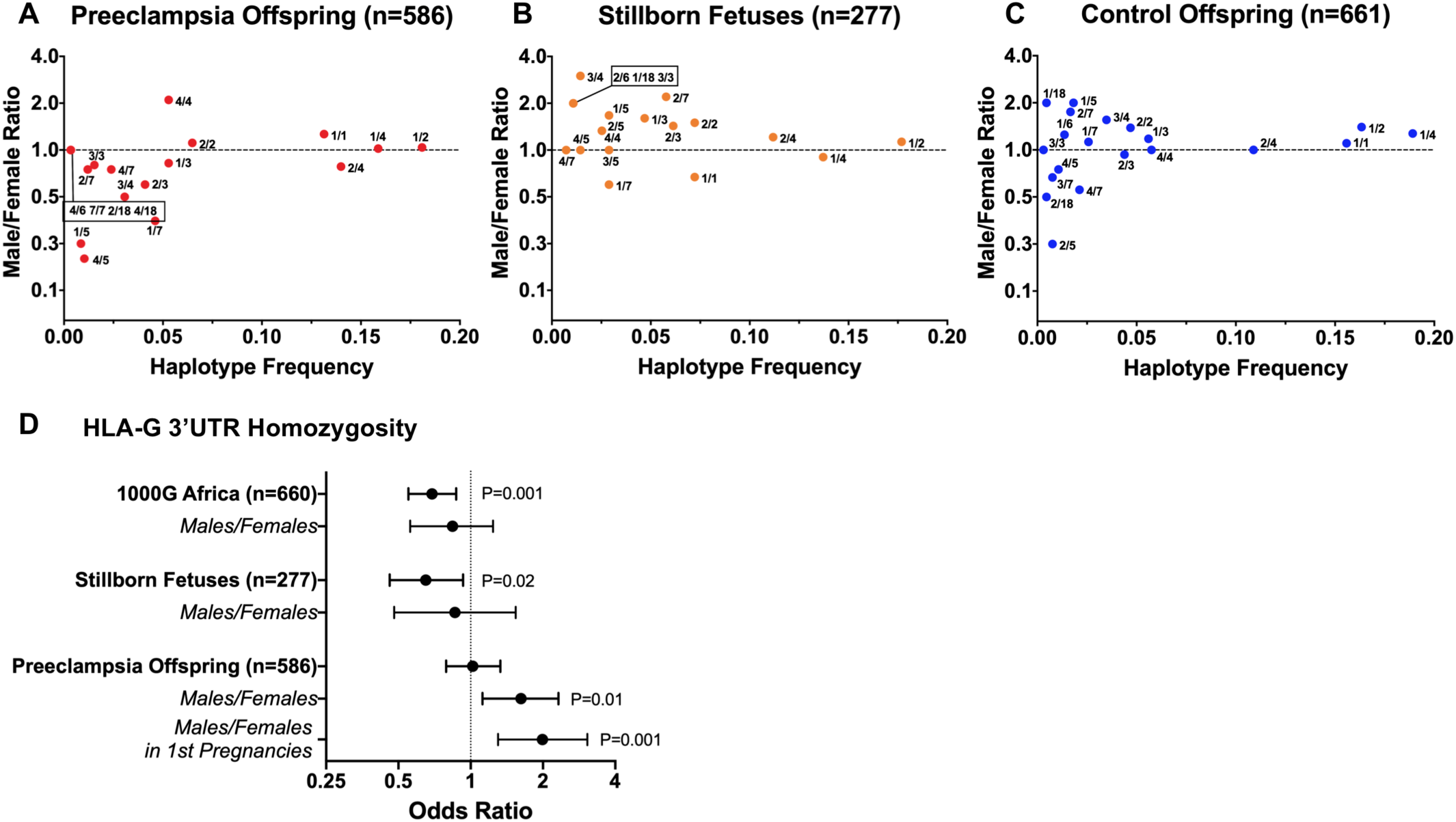
*HLA-G* 3’UTR Haplotypes, Sex Ratio and Placental Expression. The distribution of *HLA-G* 3’UTR diplotypes (e.g. 1/2 defining the combination of UTR-1 and UTR-2) and the associated birth sex ratios in (A) preeclampsia, (B) stillborn fetuses, and (C) control offspring. The FINNPEC dataset (A and C) indicates that the *HLA-G* diplotype is the main determinant of the bias in the sex ratio (GLM; allele 1: F_5,21_=0.44, P=0.82; allele 2: F_7,21_=3.51, P=0.01; allele 1 by allele 2 interaction: F_13,21_=9.15, P<0.001; haplotype frequency: F_1,21_=0.67, P=0.42; maternal condition: F_1,21_=3.58, P=0.07). Horizontal lines indicate the balanced sex ratio of 1.0. (D) The odds ratios (dots) and 95% confidence intervals (whiskers) for homozygous versus non-homozygous individuals for *HLA-G* 3’UTR. Only P values <0.05 are shown (Chi-Square test). 1000G= 1000 Genomes.

We further tested segregation of individuals homozygous for *HLA-G* 3’UTR (Fig. 2D). Decreased *HLA-G* homozygosity was observed in Africans (18.3% homozygous) versus Non-Africans (24.4%), and in the Swedish stillbirth cohort (18.4%) versus populations controls (25.7%), without sex-specific differences. Preeclamptic (26.8%) versus control offspring (26.3%) showed no differences in the overall homozygosity. However, homozygous males (31.7%; 35.2% in first pregnancies) versus females (22.3%; 21.4% in first pregnancies) were overrepresented in preeclamptic births in the FINNPEC cohort. The same association between homozygous male offspring and preeclampsia was observed in the preeclampsia family series *(16)*. Collectively, these results provide further support for balancing selection. While heterozygote advantage looks apparent for Africans, heterozygosity also shows an association with stillbirth in the Swedish cohort. In contrast, selection against *HLA-G* 3’UTR homozygous male offspring in preeclampsia supports the need for the fetus to differ from maternal HLA *(3)*, refines the locus for the findings of HLA similarity in couples with recurrent miscarriages *(4)*, and sheds light on the first-pregnancy predominance of preeclampsia *(24)*.

To support balancing selection on the HLA-G locus, we observed different frequencies of *HLA-G* 3’UTR diplotypes in offspring from preeclamptic and control pregnancies in the FINNPEC cohort: UTR-2 associated diplotypes dominated in preeclampsia (Fig. S3). The two major haplotypes, UTR-1 and UTR-2, differ for five polymorphic sites, mostly associate with the same HLA-G protein, and have global frequencies of 20-30% each *(18, 25)*. Of them, the evolutionarily most recent UTR-1 showed reduced frequency (P=0.01; OR 0.77, 95% CI: 0.63-0.94) and the ancestral haplotype UTR-2 increased frequency (P=0.01; OR 1.34, 95% CI: 1.07-1.69) in preeclamptic versus control offspring of first-time mothers. For UTR-2, the association remained when offspring from later pregnancies were included (P=0.04; OR 1.22, 95% CI: 1.01-1.47) (Table S1). The observed opposite effects of UTR-1 and UTR-2 on preeclampsia risk in first pregnancies support the advantage of divergent alleles and heterozygosity. Moreover, previous studies link UTR-2 to low expression, immune-mediated disorders, and pregnancy complications, and UTR-1 to high *HLA-G* expression *(6, 7)*. In line with this, the stillbirth cohort showed a similar but more significant protective effect of UTR-1 (P=0.0001; OR 0.65, 95% CI: 0.53-0.81). Conversely, UTR-5 was detected as a risk haplotype for stillbirth (P=0.03; OR 1.72, 95% CI: 1.06-2.79) (Table S2). UTR-5 is considered the oldest of all haplotypes and is closest to orthologous sequences in non-human primates *(25)*. We further confirmed that the 14 bp insertion polymorphism, previously implicated in pregnancy complications and present in haplotypes UTR-2, −5 and −7 *(7, 18)*, showed a modest association with both stillbirths (P=0.04; OR 1.24, 95% CI: 1.01-1.52) and preeclampsia (P=0.03; OR 1.21, 95% CI: 1.02 to 1.44).

To test the hypothesis of maternal immunoreactivity as the mechanism of fetal selection, we studied placental RNA expression of 136 genes (Fig. 3, Table S3, and Table S4), confirmed sex-linked *HLA-G* downregulation in preeclampsia (Fig. S4, Fig. S5, and S6), and found support for balancing selection (Fig. S7) *(16)*. No other immune-related genes showed compensatory effects. The marked upregulation of *interferon alpha-1* (*IFNA1*) in preeclampsia suggests that interferon signaling drives loss of fetal immunotolerance and might be targeted by its inhibitors, such as hydroxychloroquine, in clinical trials of preeclampsia and miscarriages *(16)*.

**Fig. 3.**
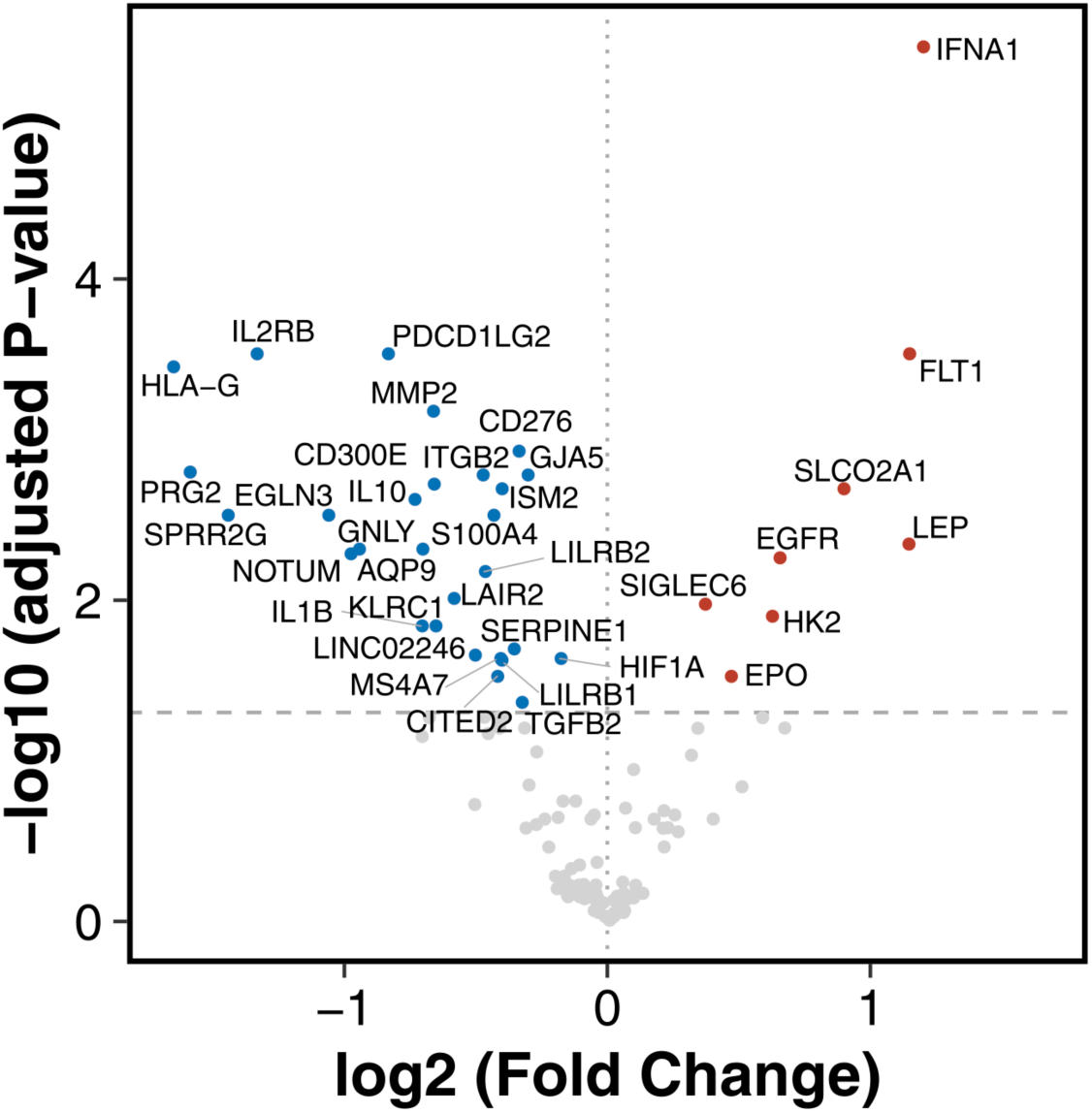
Downregulation of *HLA-G* and Upregulation of *IFNA1* in Preeclampsia. Volcano plot of downregulated (blue dots) and upregulated (red dots) genes (P_adj_<0.05) in placentas from women with severe preeclampsia (n=81) versus controls (n=63). Horizontal dashed line indicates P_adj_=0.05 and vertical dotted line indicates no fold change.

We provide the first evidence of a gene modulating human sex ratio and support *HLA-G* as a sexually antagonistic locus determining maternal-fetal interaction and pregnancy success. Furthermore, we provide empirical evidence that balancing selection is acting on contemporary human populations and complicates human pregnancies. Balancing selection of *HLA-G* haplotypes clarifies the evolutionary paradox of preeclampsia and its highest prevalence in Sub-Saharan Africa *(26)*. How can a genetic trait potentially fatal for both the fetus and the mother show heritability estimates of 50% *(27)* and persist across human populations? We propose that low *HLA-G* expression, observed also in trophoblasts infected by *Plasmodium falciparum (28)* and some viruses (CMV, HSV) *(29)*, is advantageous in pathogen-rich environments to mount protective maternal immune responses, while concurrently detrimental, causing unfavorable maternal anti-fetal immune reactions, fetal loss, and preeclampsia *(16)*. In agreement with this scenario, ancestral fetal *HLA-G* haplotypes showed associations with preeclampsia and stillbirths in our series, thereby supporting their disadvantage in modern environments. Furthermore, high *HLA-G* expression is, however, associated with higher implantation rates *(30)* and uneventful pregnancies as presented here, but might confer an increased risk of offspring to malaria and other parasitic infections *(31, 32)*. These mechanisms might be in part mediated by the *fms-like tyrosine kinase 1* (*FLT1*) gene, as our results link balancing selection of *HLA-G* regulatory haplotypes also to asymmetry for placental *FLT1* expression. Notably, high fetal *FLT1* levels confer resistance against placental malaria and fetal loss *(33)*, and maternal hypertension and elevated sFLT1 levels are not specific to preeclampsia but arise in first-time mothers with placental malaria *(34, 35)*. To support their shared pathogenesis, placental malaria and preeclampsia show high rates and seasonal co-occurrence in Africa *(36)*, where both the *HLA-G* low-expressing “null allele” and high-expressing *FLT1* alleles are under positive selection *(33, 37)*, and the frequency and sex ratios associated with *HLA-G* 3’UTR haplotypes differ from other populations as shown here.

The majority of humans are HLA heterozygous for the need to provide more productive immune responses to pathogens *(3, 21)*. While *HLA-G* shows physiological expression almost uniquely in trophoblasts, its high expression that benefits the viability of fetuses during pregnancy may trade-off with *HLA-G* neoexpression, promoting immune evasion of viruses and malaria as well as tumors later in life *(6, 38)*. Therefore, the benefit of differential *HLA-G* expression may be dependent on the time and place, pathogens, and the tissue of expression, providing a role for balancing selection on immune-related traits beyond pregnancy in human evolution.

## Acknowledgments

We thank all the FINNPEC study participants and the investigators of the FINNPEC study group (listed in Supplementary Appendix). We appreciate the expert technical assistance of Auli Saarinen, Eija Kortelainen, and Susanna Mehtälä and contribution of the assisting personnel of the FINNPEC Study. Biomedicum Functional Genomics Unit (FuGU), University of Helsinki, is acknowledged for STRT library sequencing services.

## Funding

S.W. received funding from the Finnish Medical Foundation, and Päivikki and Sakari Sohlberg Foundation. M.Y. was supported by the Karolinska Institutet Research Foundation, Scandinavia-Japan Sasakawa Foundation, Japan Eye Bank Association, Astellas Foundation for Research on Metabolic Disorders, and Japan Society for the Promotion of Science Overseas Research Fellowships. Work in the J.K. laboratory is supported by Knut and Alice Wallenberg Foundation (KAW 2015.0096), Swedish Research Council, Medical Society Liv och Hälsa (Life and Health), and Sigrid Jusélius Foundation. J.K. is a recipient of the Wolfson Research Merit Award by the Royal Society. The FINNPEC study has received funding from the Competitive State Research Financing of the Expert Responsibility area of Helsinki University Hospital (TYH2018305), Jane and Aatos Erkko Foundation, Päivikki and Sakari Sohlberg Foundation, Academy of Finland (grants 121196, 134957 and 278941), Research Funds of the University of Helsinki, Finnish Medical Foundation, Finska Läkaresällskapet, Novo Nordisk Foundation, Finnish Foundation for Pediatric Research, Emil Aaltonen Foundation, and Sigrid Jusélius Foundation.

## Author contributions

S.W. and J.K. conceived and designed the study. E.S., E.I., K.P., E.K., S.H., H.L. and J.K. collected the patient cohorts and prepared the samples. S.W., H.S., I.M.H., S.E., T.S. and K.K. performed the experiments. S.W., M.Y., H.T., M.G., S.K., J.W., N.L., J.L., M.M. and J.K. generated, analyzed, and interpreted the data. S.W., M.Y. and J.K. wrote the manuscript and all authors substantially revised the manuscript.

## Competing interests

Authors declare no competing interests.

## Data and materials availability

Gene expression data were deposited in NCBI’s Gene Expression Omnibus and are accessible through the accession GSE125460 (https://www.ncbi.nlm.nih.gov/geo/query/acc.cgi?acc=GSE125460). All other data is available in the main text or the supplementary materials.

## Supplementary Materials

Materials and Methods

Supplementary Text

Figures S1-S19

Tables S1-S6

References (39-137)

## References and Notes

1. S. H. Orzack et al., The human sex ratio from conception to birth. Proc. Natl. Acad. Sci. U. S. A. 112, E2102–11 (2015).

2. R. Fisher, The genetical theory of natural selection (Clarendon Press, Oxford, 1930).

3. F. L. Black, P. W. Hedrick, Strong balancing selection at HLA loci: evidence from segregation in South Amerindian families. Proc. Natl. Acad. Sci. U. S. A. 94, 12452–12456 (1997).

4. C. Ober, T. Hyslop, S. Elias, L. R. Weitkamp, W. W. Hauck, Human leukocyte antigen matching and fetal loss: results of a 10 year prospective study. Hum. Reprod. 13, 33–38 (1998).

5. J. Pollheimer, S. Vondra, J. Baltayeva, A. G. Beristain, M. Knofler, Regulation of Placental Extravillous Trophoblasts by the Maternal Uterine Environment. Front. Immunol. 9, 2597 (2018).

6. E. D. Carosella, N. Rouas-Freiss, D. Tronik-Le Roux, P. Moreau, J. LeMaoult, HLA-G: An Immune Checkpoint Molecule. Adv. Immunol. 127, 33–144 (2015).

7. L. Lynge Nilsson, S. Djurisic, T. V. Hviid, Controlling the Immunological Crosstalk during Conception and Pregnancy: HLA-G in Reproduction. Front. Immunol. 5, 198 (2014).

8. S. Djurisic, T. V. Hviid, HLA Class Ib Molecules and Immune Cells in Pregnancy and Preeclampsia. Front. Immunol. 5, 652 (2014).

9. I. L. Sargent, A. M. Borzychowski, C. W. Redman, NK cells and pre-eclampsia. J. Reprod. Immunol. 76, 40–44 (2007).

10. C. W. Redman, I. L. Sargent, Immunology of pre-eclampsia. Am. J. Reprod. Immunol. 63, 534–543 (2010).

11. D. Jaskolka, R. Retnakaran, B. Zinman, C. K. Kramer, Fetal sex and maternal risk of pre-eclampsia/eclampsia: a systematic review and meta-analysis. Bjog. 124, 553–560 (2017).

12. R. Arngrimsson, J. J. Walker, R. T. Geirsson, S. Bjornsson, A low male/female sex ratio in offspring of women with a family history of pre-eclampsia and eclampsia. Br. J. Obstet. Gynaecol. 100, 496–497 (1993).

13. L. J. Vatten, R. Skjaerven, Offspring sex and pregnancy outcome by length of gestation. Early Hum. Dev. 76, 47–54 (2004).

14. Global Pregnancy Collaboration: et al., Fetal sex-specific differences in gestational age at delivery in pre-eclampsia: a meta-analysis. Int. J. Epidemiol. 46, 632–642 (2017).

15. D. W. Cooper, J. A. Hill, L. C. Chesley, C. I. Bryans, Genetic control of susceptibility to eclampsia and miscarriage. Br. J. Obstet. Gynaecol. 95, 644–653 (1988).

16. See supplementary materials.

17. U. Muller, H-Y antigens. Hum. Genet. 97, 701–704 (1996).

18. E. C. Castelli et al., Insights into HLA-G Genetics Provided by Worldwide Haplotype Diversity. Front. Immunol. 5, 476 (2014).

19. 1000 Genomes Project Consortium et al., A global reference for human genetic variation. Nature. 526, 68–74 (2015).

20. A. Nakimuli et al., Pregnancy, parturition and preeclampsia in women of African ancestry. Am. J. Obstet. Gynecol. 210, 510-520.e1 (2014).

21. P. Parham, T. Ohta, Population biology of antigen presentation by MHC class I molecules. Science. 272, 67–74 (1996).

22. T. L. Lenz, Adaptive value of novel MHC immune gene variants. Proc. Natl. Acad. Sci. U. S. A. 115, 1414–1416 (2018).

23. K. P. Phillips et al., Immunogenetic novelty confers a selective advantage in host-pathogen coevolution. Proc. Natl. Acad. Sci. U. S. A. 115, 1552–1557 (2018).

24. E. A. Steegers, P. von Dadelszen, J. J. Duvekot, R. Pijnenborg, Pre-eclampsia. Lancet. 376, 631–644 (2010).

25. A. Sabbagh et al., Worldwide genetic variation at the 3’ untranslated region of the HLA-G gene: balancing selection influencing genetic diversity. Genes Immun. 15, 95–106 (2014).

26. L. Say et al., Global causes of maternal death: a WHO systematic analysis. Lancet Glob. Health. 2, e323–33 (2014).

27. S. Cnattingius, M. Reilly, Y. Pawitan, P. Lichtenstein, Maternal and fetal genetic factors account for most of familial aggregation of preeclampsia: a population-based Swedish cohort study. Am. J. Med. Genet. A. 130A, 365–371 (2004).

28. H. Sartelet et al., Less HLA-G expression in Plasmodium falciparum-infected third trimester placentas is associated with more natural killer cells. Placenta. 26, 505–511 (2005).

29. K. Racicot, G. Mor, Risks associated with viral infections during pregnancy. J. Clin. Invest. 127, 1591–1599 (2017).

30. V. Rebmann, M. Switala, I. Eue, H. Grosse-Wilde, Soluble HLA-G is an independent factor for the prediction of pregnancy outcome after ART: a German multi-centre study. Hum. Reprod. 25, 1691–1698 (2010).

31. A. Sabbagh et al., The role of HLA-G in parasitic diseases. Hla. 91, 255–270 (2018).

32. T. C. d’Almeida et al., High level of soluble human leukocyte antigen (HLA)-G at beginning of pregnancy as predictor of risk of malaria during infancy. Sci. Rep. 9, 9160-019-45688-w (2019).

33. A. Muehlenbachs, M. Fried, J. Lachowitzer, T. K. Mutabingwa, P. E. Duffy, Natural selection of FLT1 alleles and their association with malaria resistance in utero. Proc. Natl. Acad. Sci. U. S. A. 105, 14488–14491 (2008).

34. S. E. Maynard et al., Excess placental soluble fms-like tyrosine kinase 1 (sFlt1) may contribute to endothelial dysfunction, hypertension, and proteinuria in preeclampsia. J. Clin. Invest. 111, 649–658 (2003).

35. A. Muehlenbachs, T. K. Mutabingwa, S. Edmonds, M. Fried, P. E. Duffy, Hypertension and maternal-fetal conflict during placental malaria. PLoS Med. 3, e446 (2006).

36. H. Sartelet, C. Rogier, I. Milko-Sartelet, G. Angel, G. Michel, Malaria associated pre-eclampsia in Senegal. Lancet. 347, 1121 (1996).

37. C. Aldrich, C. Wambebe, L. Odama, A. Di Rienzo, C. Ober, Linkage disequilibrium and age estimates of a deletion polymorphism (1597DeltaC) in HLA-G suggest non-neutral evolution. Hum. Immunol. 63, 405–412 (2002).

38. L. Amiot, N. Vu, M. Samson, Immunomodulatory properties of HLA-G in infectious diseases. J. Immunol. Res. 2014, 298569 (2014).

39. D. R. Zerbino et al., Ensembl 2018. Nucleic Acids Res. 46, D754–D761 (2018).

40. E. Sahlin et al., Identification of putative pathogenic single nucleotide variants (SNVs) in genes associated with heart disease in 290 cases of stillbirth. PLoS One. 14, e0210017 (2019).

41. K. H. Cui, G. M. Warnes, R. Jeffrey, C. D. Matthews, Sex determination of preimplantation embryos by human testis-determining-gene amplification. Lancet. 343, 79–82 (1994).

42. C. Gunter et al., Re-examination of factors associated with expansion of CGG repeats using a single nucleotide polymorphism in FMR1. Hum. Mol. Genet. 7, 1935–1946 (1998).

43. T. Jaaskelainen et al., Cohort profile: the Finnish Genetics of Pre-eclampsia Consortium (FINNPEC). BMJ Open. 6, e013148-2016-013148 (2016).

44. E. C. Castelli et al., The genetic structure of 3’untranslated region of the HLA-G gene: polymorphisms and haplotypes. Genes Immun. 11, 134–141 (2010).

45. M. Gauster, G. Moser, K. Orendi, B. Huppertz, Factors involved in regulating trophoblast fusion: potential role in the development of preeclampsia. Placenta. 30 Suppl A, S49–54 (2009).

46. R. Apps et al., Genome-wide expression profile of first trimester villous and extravillous human trophoblast cells. Placenta. 32, 33–43 (2011).

47. C. Q. Lee et al., What Is Trophoblast? A Combination of Criteria Define Human First-Trimester Trophoblast. Stem Cell. Reports. 6, 257–272 (2016).

48. FANTOM Consortium and the RIKEN PMI and CLST (DGT) et al., A promoter-level mammalian expression atlas. Nature. 507, 462–470 (2014).

49. M. G. Petroff, A. Perchellet, B7 family molecules as regulators of the maternal immune system in pregnancy. Am. J. Reprod. Immunol. 63, 506–519 (2010).

50. D. Monk, Genomic imprinting in the human placenta. Am. J. Obstet. Gynecol. 213, S152–62 (2015).

51. Y. Zhou, C. H. Damsky, K. Chiu, J. M. Roberts, S. J. Fisher, Preeclampsia is associated with abnormal expression of adhesion molecules by invasive cytotrophoblasts. J. Clin. Invest. 91, 950–960 (1993).

52. R. W. Redline, P. Patterson, Pre-eclampsia is associated with an excess of proliferative immature intermediate trophoblast. Hum. Pathol. 26, 594–600 (1995).

53. V. D. Winn et al., Severe preeclampsia-related changes in gene expression at the maternal-fetal interface include sialic acid-binding immunoglobulin-like lectin-6 and pappalysin-2. Endocrinology. 150, 452–462 (2009).

54. Y. Zhou et al., Reversal of gene dysregulation in cultured cytotrophoblasts reveals possible causes of preeclampsia. J. Clin. Invest. 123, 2862–2872 (2013).

55. C. W. Redman, I. L. Sargent, A. C. Staff, IFPA Senior Award Lecture: making sense of pre-eclampsia - two placental causes of preeclampsia? Placenta. 35 Suppl, S20–5 (2014).

56. T. Kaartokallio et al., Gene expression profiling of pre-eclamptic placentae by RNA sequencing. Sci. Rep. 5, 14107 (2015).

57. S. F. Altschul, W. Gish, W. Miller, E. W. Myers, D. J. Lipman, Basic local alignment search tool. J. Mol. Biol. 215, 403–410 (1990).

58. H. Teder et al., TAC-seq: targeted DNA and RNA sequencing for precise biomarker molecule counting. NPJ Genom. Med. 3, 34-018-0072-5. eCollection 2018 (2018).

59. R. Edgar, M. Domrachev, A. E. Lash, Gene Expression Omnibus: NCBI gene expression and hybridization array data repository. Nucleic Acids Res. 30, 207–210 (2002).

60. P. Ewels, M. Magnusson, S. Lundin, M. Kaller, MultiQC: summarize analysis results for multiple tools and samples in a single report. Bioinformatics. 32, 3047–3048 (2016).

61. T. Kivioja et al., Counting absolute numbers of molecules using unique molecular identifiers. Nat. Methods. 9, 72–74 (2011).

62. M. I. Love, W. Huber, S. Anders, Moderated estimation of fold change and dispersion for RNA-seq data with DESeq2. Genome Biol. 15, 550-014-0550-8 (2014).

63. M. E. Ritchie et al., limma powers differential expression analyses for RNA-sequencing and microarray studies. Nucleic Acids Res. 43, e47 (2015).

64. X. Robin et al., pROC: an open-source package for R and S+ to analyze and compare ROC curves. BMC Bioinformatics. 12, 77-2105-12-77 (2011).

65. S. Islam et al., Characterization of the single-cell transcriptional landscape by highly multiplex RNA-seq. Genome Res. 21, 1160–1167 (2011).

66. S. Islam et al., Quantitative single-cell RNA-seq with unique molecular identifiers. Nat. Methods. 11, 163–166 (2014).

67. K. Krjutskov et al., Globin mRNA reduction for whole-blood transcriptome sequencing. Sci. Rep. 6, 31584 (2016).

68. K. Krjutskov et al., Single-cell transcriptome analysis of endometrial tissue. Hum. Reprod. 31, 844–853 (2016).

69. S. Katayama, V. Tohonen, S. Linnarsson, J. Kere, SAMstrt: statistical test for differential expression in single-cell transcriptome with spike-in normalization. Bioinformatics. 29, 2943–2945 (2013).

70. D. Kim, J. M. Paggi, C. Park, C. Bennett, S. L. Salzberg, Graph-based genome alignment and genotyping with HISAT2 and HISAT-genotype. Nat. Biotechnol. 37, 907–915 (2019).

71. Y. Liao, G. K. Smyth, W. Shi, featureCounts: an efficient general purpose program for assigning sequence reads to genomic features. Bioinformatics. 30, 923–930 (2014).

72. S. Katayama et al., Guide for library design and bias correction for large-scale transcriptome studies using highly multiplexed RNAseq methods. BMC Bioinformatics. 20, 418-019-3017-9 (2019).

73. R. Apps, L. Gardner, A. Moffett, A critical look at HLA-G. Trends Immunol. 29, 313–321 (2008).

74. Z. Awamleh, G. B. Gloor, V. K. M. Han, Placental microRNAs in pregnancies with early onset intrauterine growth restriction and preeclampsia: potential impact on gene expression and pathophysiology. BMC Med. Genomics. 12, 91-019-0548-x (2019).

75. H. Laivuori et al., Susceptibility loci for preeclampsia on chromosomes 2p25 and 9p13 in Finnish families. Am. J. Hum. Genet. 72, 168–177 (2003).

76. E. W. Triche, K. K. Harland, E. H. Field, L. M. Rubenstein, A. F. Saftlas, Maternal-fetal HLA sharing and preeclampsia: variation in effects by seminal fluid exposure in a case-control study of nulliparous women in Iowa. J. Reprod. Immunol. 101-102, 111–119 (2014).

77. S. B. Cheng, S. Sharma, Interleukin-10: a pleiotropic regulator in pregnancy. Am. J. Reprod. Immunol. 73, 487–500 (2015).

78. H. P. Moll, T. Maier, A. Zommer, T. Lavoie, C. Brostjan, The differential activity of interferon-alpha subtypes is consistent among distinct target genes and cell types. Cytokine. 53, 52–59 (2011).

79. L. B. Ivashkiv, L. T. Donlin, Regulation of type I interferon responses. Nat. Rev. Immunol. 14, 36–49 (2014).

80. R. J. Levine et al., Circulating angiogenic factors and the risk of preeclampsia. N. Engl. J. Med. 350, 672–683 (2004).

81. A. L. Bayer, A. Pugliese, T. R. Malek, The IL-2/IL-2R system: from basic science to therapeutic applications to enhance immune regulation. Immunol. Res. 57, 197–209 (2013).

82. S. J. Fisher, Why is placentation abnormal in preeclampsia? Am. J. Obstet. Gynecol. 213, S115–22 (2015).

83. A. Waysbort et al., Experimental study of transplacental passage of alpha interferon by two assay techniques. Antimicrob. Agents Chemother. 37, 1232–1237 (1993).

84. P. Magnus, A. Eskild, Seasonal variation in the occurrence of pre-eclampsia. Bjog. 108, 1116–1119 (2001).

85. D. Fisman, Seasonality of viral infections: mechanisms and unknowns. Clin. Microbiol. Infect. 18, 946–954 (2012).

86. M. Cappelletti et al., Type I interferons regulate susceptibility to inflammation-induced preterm birth. JCI Insight. 2, e91288 (2017).

87. G. Engels et al., Pregnancy-Related Immune Adaptation Promotes the Emergence of Highly Virulent H1N1 Influenza Virus Strains in Allogenically Pregnant Mice. Cell. Host Microbe. 21, 321–333 (2017).

88. K. Racicot et al., Cutting Edge: Fetal/Placental Type I IFN Can Affect Maternal Survival and Fetal Viral Load during Viral Infection. J. Immunol. 198, 3029–3032 (2017).

89. L. J. Yockey et al., Type I interferons instigate fetal demise after Zika virus infection. Sci. Immunol. 3, 10.1126/sciimmunol.aao1680 (2018).

90. M. K. Crow, Advances in understanding the role of type I interferons in systemic lupus erythematosus. Curr. Opin. Rheumatol. 26, 467–474 (2014).

91. L. Zitvogel, L. Galluzzi, O. Kepp, M. J. Smyth, G. Kroemer, Type I interferons in anticancer immunity. Nat. Rev. Immunol. 15, 405–414 (2015).

92. V. E. Reyes, G. R. Klimpel, Interferon alpha/beta synthesis during acute graft-versus-host disease. Transplantation. 43, 412–416 (1987).

93. M. G. Cleveland, R. G. Lane, G. R. Klimpel, Enhanced interferon-alpha/beta (IFN-alpha/beta) and defective IFN-gamma production in chronic graft versus host disease: a potential mechanism for immunosuppression. Cell. Immunol. 110, 120–130 (1987).

94. M. L. Alegre et al., Antagonistic effect of toll-like receptor signaling and bacterial infections on transplantation tolerance. Transplantation. 87, S77–9 (2009).

95. M. Matz et al., The regulation of interferon type I pathway-related genes RSAD2 and ETV7 specifically indicates antibody-mediated rejection after kidney transplantation. Clin. Transplant. 32, e13429 (2018).

96. M. E. Clowse, M. Jamison, E. Myers, A. H. James, A national study of the complications of lupus in pregnancy. Am. J. Obstet. Gynecol. 199, 127.e1-127.e6 (2008).

97. S. Hong et al., Longitudinal profiling of human blood transcriptome in healthy and lupus pregnancy. J. Exp. Med. 216, 1154–1169 (2019).

98. K. Sacre, L. A. Criswell, J. M. McCune, Hydroxychloroquine is associated with impaired interferon-alpha and tumor necrosis factor-alpha production by plasmacytoid dendritic cells in systemic lupus erythematosus. Arthritis Res. Ther. 14, R155 (2012).

99. M. Leroux et al., Impact of hydroxychloroquine on preterm delivery and intrauterine growth restriction in pregnant women with systemic lupus erythematosus: a descriptive cohort study. Lupus. 24, 1384–1391 (2015).

100. M. R. Seo et al., Hydroxychloroquine treatment during pregnancy in lupus patients is associated with lower risk of preeclampsia. Lupus. 28, 722–730 (2019).

101. W. Marder, Update on pregnancy complications in systemic lupus erythematosus. Curr. Opin. Rheumatol. 31, 650–658 (2019).

102. T. Wang, X. Z. Shi, W. H. Wu, Crosstalk analysis of dysregulated pathways in preeclampsia. Exp. Ther. Med. 17, 2298–2304 (2019).

103. R. Rahman et al., The effects of hydroxychloroquine on endothelial dysfunction. Pregnancy Hypertens. 6, 259–262 (2016).

104. M. Gomez-Guzman et al., Chronic hydroxychloroquine improves endothelial dysfunction and protects kidney in a mouse model of systemic lupus erythematosus. Hypertension. 64, 330–337 (2014).

105. A. Scharfe-Nugent et al., TLR9 provokes inflammation in response to fetal DNA: mechanism for fetal loss in preterm birth and preeclampsia. J. Immunol. 188, 5706–5712 (2012).

106. B. Cao, L. A. Parnell, M. S. Diamond, I. U. Mysorekar, Inhibition of autophagy limits vertical transmission of Zika virus in pregnant mice. J. Exp. Med. 214, 2303–2313 (2017).

107. D. J. Schust, A. B. Hill, H. L. Ploegh, Herpes simplex virus blocks intracellular transport of HLA-G in placentally derived human cells. J. Immunol. 157, 3375–3380 (1996).

108. Y. Jun et al., Human cytomegalovirus gene products US3 and US6 down-regulate trophoblast class I MHC molecules. J. Immunol. 164, 805–811 (2000).

109. F. Arechavaleta-Velasco, H. Koi, J. F. Strauss 3rd, S. Parry, Viral infection of the trophoblast: time to take a serious look at its role in abnormal implantation and placentation? J. Reprod. Immunol. 55, 113–121 (2002).

110. F. McNab, K. Mayer-Barber, A. Sher, A. Wack, A. O’Garra, Type I interferons in infectious disease. Nat. Rev. Immunol. 15, 87–103 (2015).

111. M. C. de Goffau et al., Human placenta has no microbiome but can contain potential pathogens. Nature. 572, 329–334 (2019).

112. A. M. Cotter, C. M. Martin, J. J. O’leary, S. F. Daly, Increased fetal DNA in the maternal circulation in early pregnancy is associated with an increased risk of preeclampsia. Am. J. Obstet. Gynecol. 191, 515–520 (2004).

113. C. Nathan, A. Ding, Nonresolving inflammation. Cell. 140, 871–882 (2010).

114. D. Andrade et al., Interferon-alpha and angiogenic dysregulation in pregnant lupus patients who develop preeclampsia. Arthritis Rheumatol. 67, 977–987 (2015).

115. W. C. HuangFu, J. Liu, R. N. Harty, S. Y. Fuchs, Cigarette smoking products suppress anti-viral effects of Type I interferon via phosphorylation-dependent downregulation of its receptor. FEBS Lett. 582, 3206–3210 (2008).

116. S. A. Karumanchi, R. J. Levine, How does smoking reduce the risk of preeclampsia? Hypertension. 55, 1100–1101 (2010).

117. S. Kovats et al., A class I antigen, HLA-G, expressed in human trophoblasts. Science. 248, 220–223 (1990).

118. M. Dahl, S. Djurisic, T. V. Hviid, The many faces of human leukocyte antigen-G: relevance to the fate of pregnancy. J. Immunol. Res. 2014, 591489 (2014).

119. C. Solier et al., Secretion of pro-apoptotic intron 4-retaining soluble HLA-G1 by human villous trophoblast. Eur. J. Immunol. 32, 3576–3586 (2002).

120. P. J. Morales, J. L. Pace, J. S. Platt, D. K. Langat, J. S. Hunt, Synthesis of beta(2)-microglobulin-free, disulphide-linked HLA-G5 homodimers in human placental villous cytotrophoblast cells. Immunology. 122, 179–188 (2007).

121. H. Kumar, T. Kawai, S. Akira, Pathogen recognition by the innate immune system. Int. Rev. Immunol. 30, 16–34 (2011).

122. G. Mitchell, R. R. Isberg, Innate Immunity to Intracellular Pathogens: Balancing Microbial Elimination and Inflammation. Cell. Host Microbe. 22, 166–175 (2017).

123. E. DiFederico, O. Genbacev, S. J. Fisher, Preeclampsia is associated with widespread apoptosis of placental cytotrophoblasts within the uterine wall. Am. J. Pathol. 155, 293–301 (1999).

124. I. P. Crocker, S. Cooper, S. C. Ong, P. N. Baker, Differences in apoptotic susceptibility of cytotrophoblasts and syncytiotrophoblasts in normal pregnancy to those complicated with preeclampsia and intrauterine growth restriction. Am. J. Pathol. 162, 637–643 (2003).

125. M. S. Longtine, B. Chen, A. O. Odibo, Y. Zhong, D. M. Nelson, Villous trophoblast apoptosis is elevated and restricted to cytotrophoblasts in pregnancies complicated by preeclampsia, IUGR, or preeclampsia with IUGR. Placenta. 33, 352–359 (2012).

126. D. Goswami et al., Excess syncytiotrophoblast microparticle shedding is a feature of early-onset pre-eclampsia, but not normotensive intrauterine growth restriction. Placenta. 27, 56–61 (2006).

127. A. M. Vigario et al., Inhibition of Plasmodium yoelii blood-stage malaria by interferon alpha through the inhibition of the production of its target cell, the reticulocyte. Blood. 97, 3966–3971 (2001).

128. I. Sebina, A. Haque, Effects of type I interferons in malaria. Immunology. 155, 176–185 (2018).

129. J. M. Gonzalez-Navajas, J. Lee, M. David, E. Raz, Immunomodulatory functions of type I interferons. Nat. Rev. Immunol. 12, 125–135 (2012).

130. Z. von Marschall et al., Effects of interferon alpha on vascular endothelial growth factor gene transcription and tumor angiogenesis. J. Natl. Cancer Inst. 95, 437–448 (2003).

131. S. Ahmad, A. Ahmed, Elevated placental soluble vascular endothelial growth factor receptor-1 inhibits angiogenesis in preeclampsia. Circ. Res. 95, 884–891 (2004).

132. C. T. Taylor, S. P. Colgan, Regulation of immunity and inflammation by hypoxia in immunological niches. Nat. Rev. Immunol. 17, 774–785 (2017).

133. P. S. Macklin, J. McAuliffe, C. W. Pugh, A. Yamamoto, Hypoxia and HIF pathway in cancer and the placenta. Placenta. 56, 8–13 (2017).

134. J. Hanna et al., Decidual NK cells regulate key developmental processes at the human fetal-maternal interface. Nat. Med. 12, 1065–1074 (2006).

135. X. Fan et al., Endometrial VEGF induces placental sFLT1 and leads to pregnancy complications. J. Clin. Invest. 124, 4941–4952 (2014).

136. K. C. Wheeler et al., VEGF may contribute to macrophage recruitment and M2 polarization in the decidua. PLoS One. 13, e0191040 (2018).

137. S. L. Adamson, sFLT1 in preeclampsia: trophoblast defense against a decidual VEGFA barrage? J. Clin. Invest. 124, 4690–4692 (2014).

